# Flexible EMG arrays with integrated electronics for scalable electrode density

**DOI:** 10.1101/2024.07.02.601782

**Authors:** Philip Anschutz, Muneeb Zia, Jiaao Lu, Matthew Williams, Amanda Jacob, Samuel Sober, Muhannad Bakir

**Affiliations:** Georgia Institute of Technology; Emory University & Georgia Institute of Technology; Emory University; Georgia Institue of Technology

**Keywords:** Polyimide, Chip integration, Wirebonding, Electrophysiology

## Abstract

Recent developments in electrode technology have demonstrated the power of flexible microelectrode arrays (FMEAs) for measuring muscle activity at high resolution. We recently introduced the Myomatrix array, an FMEA optimized for measuring the activity of individual motor units (the collection of muscle fibers innervated by a single motor neuron) [1] in freely behaving animals. Although FMEAs are fundamentally changing the way electromyography (EMG) is acquired, the number of recording channels is limited by the size of the plug that interfaces with the signal processing hardware and the density of electrode connections on the array. Increasing EMG channel count and supporting electrophysiological studies in smaller animals depends on two seemingly incompatible goals: reducing device size while increasing the number of recording channels. The solution to these goals is to increase the channel count per wire output. Current off-the-shelf designs require a separate headstage and FMEA to be used simultaneously. In our prior devices [1], each FMEA had a dedicated wire output for every electrode input, creating a channel density of 1:1. To improve this channel density, we have developed an FMEA with an integrated digital amplifier (Bare-Die RHD2216, INTAN INC, USA). The design of the FMEA reduces the footprint of the device’s back end by 74% and relocates the Intan bare die from the headstage to the FMEA itself, creating a channel density of 1:3.2. Our methodology combines standard FMEA microfabrication with wire-bonding and surface-mounted components, enabling direct integration into a Serial Peripheral Interface (SPI) connection into the device itself, without any separate headstage. With this initial device there is a 1 : 3.2 channel density; however, our method allows using other bare die amplifiers (Intan, Inc., USA) for a channel density of 1:12.8. Our findings present a robust technique for chip embedding in custom FMEAs applicable to in-vivo electrophysiology

## I. Introduction

### A. The need for Application Specific EMG Devices

FMEAs have emerged as a new tool in measuring high signal-to-noise ratio (SNR) electromyography (EMG) applications. Customizing the electrode shape to fit a specific muscle target ensures a precise fit and optimally positions the electrodes for recording. These devices measure the electric differential caused by the contraction of the muscle fibers. We recently introduced the Myomatrix array, an FMEA optimized for measuring the activity of individual motor units [1]in freely behaving animals.

While neural recording devices have advanced rapidly over the past decade, high resolution EMG devices have not seen the same advances in custom hardware that neural devices have experienced [2], [3]. While measuring neuron activity is one approach to behavioral neuroscience, measuring the output in EMG is another approach. Combining these approaches can also be a powerful tool in diagnosing neuromuscular disorders, like Parkinson’s [4], ALS [5], and Alzheimer’s disease [6]. Clinically there is a need for quantitative biomarkers for motor diseases. An example of this is the absence of a quantitative test for Parkinson’s disease that can say if it is afflicting an individual [7]. With the Myomatrix arrays high SNR EMG can be acquired. There is a need to improve upon these devices to enable future studies in neuro-muscular disorders and discover key biomarkers that could aid in disease discovery and treatment [8].

### B. Approaching Device Architecture

Recently, there has been more interest in tailored applications for FMEAs that require smaller footprint devices and increasing the number of electrodes that are present on the device [9][10][11]. Currently we are not limited by electrode size requirements, meaning we can fabricate smaller electrodes on a smaller footprint FMEA. The limiting factor is the hardware requirements that would be needed to communicate with 10 times or 100 times the number of electrodes that are currently found in FMEA’s. The solution to this is to increase channel density. In our prior devices [1], each FMEA had a dedicated wire output for every electrode input, creating a channel density of 1:1. This design choice creates a bottleneck for increasing electrode count, and shrinking the device connections. Higher electrode counts allow for a higher resolution of the signal as the ions are released and the muscle contracts. By increasing this resolution we can find new biomarkers that are not apparent with larger, fewer electrodes [12].

To create a device that is best suited for EMG applications the analog signal from the electrodes needs to be converted to a digital output on the device, and the device itself needs to be small enough to fit the animal model it is intended for. Moving the analog-to-digital conversion closer to the signal source also reduces noise. Another consideration is that when scaling this device, we want to make it as user friendly as possible, and be able to plug into the RHD headstage amplifier board. For this reason, we have chosen to work with the INTAN RHD amplifier chip (INTAN INC, USA). The specific chip chosen is the 2216, because it records from 32 bipolar (differential) channels which is well suited for long-term EMG recording. For the Myomatrix design, we chose to work with a 32-channel array designed for a mouse forelimb recording from our previous work [1]. Our approach is to build a robust implantation process that can be modified to the many different animal models that are used in EMG research. For our first device we chose a mouse model, and created a render of the device to see if the size requirements of the embedded electronics themselves created a large advantage over the INTAN headstage. This rendering is shown in Figure 1. In this work, we propose a FMEA with an embedded INTAN 2216, to reduce the footprint of the device and increase channel density.

**Fig. 1.**
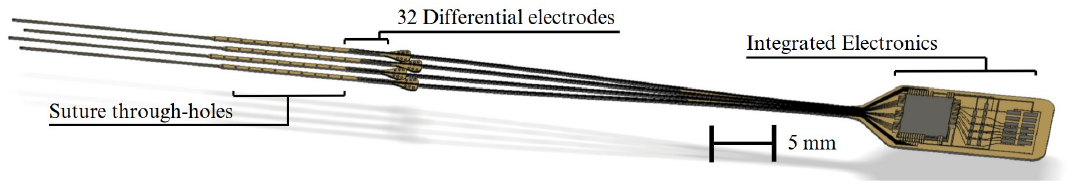
This render was used to see how small we could shrink the backend footprint of the device, and to see the size advantage it would have over the commercially available INTAN headstage

## II. Methods

### A. Design Resources

We first took the RHD 2216 headstage design and applied it to the backend of the 32-channel array designed for a mouse forelimb recording [1]. The design was constructed by taking the resources made available on INTAN’s website [13] and recreating the connections on our device.

### B. Microfabrication

To begin, 15µm of Polyimide (Pyralin 2611, HD Microsystems, USA) was spin-coated onto a 4” silicon handle wafer and cured in a nitrogen environment. After curing, a metal liftoff (20nm Ti, 300nm Au) formed the traces for the device. Following liftoff, VM652 was spin-coated on the entire wafer, and then the top Polyimide was spin-coated to 3µm. After curing, a positive resist was patterned to make an etch mask. A reactive ion etch opens the electrodes and the connections for the surface mount components and the wirebonding pads. Next we thickened the wirebonding pads so that they are compatible with commercial wirebonders. To achieve this compatibility, another liftoff process with 300nm Cu and 20nm Au is deposited on the open pads on the backend of the device as seen in Figure 2.

**Fig. 2.**
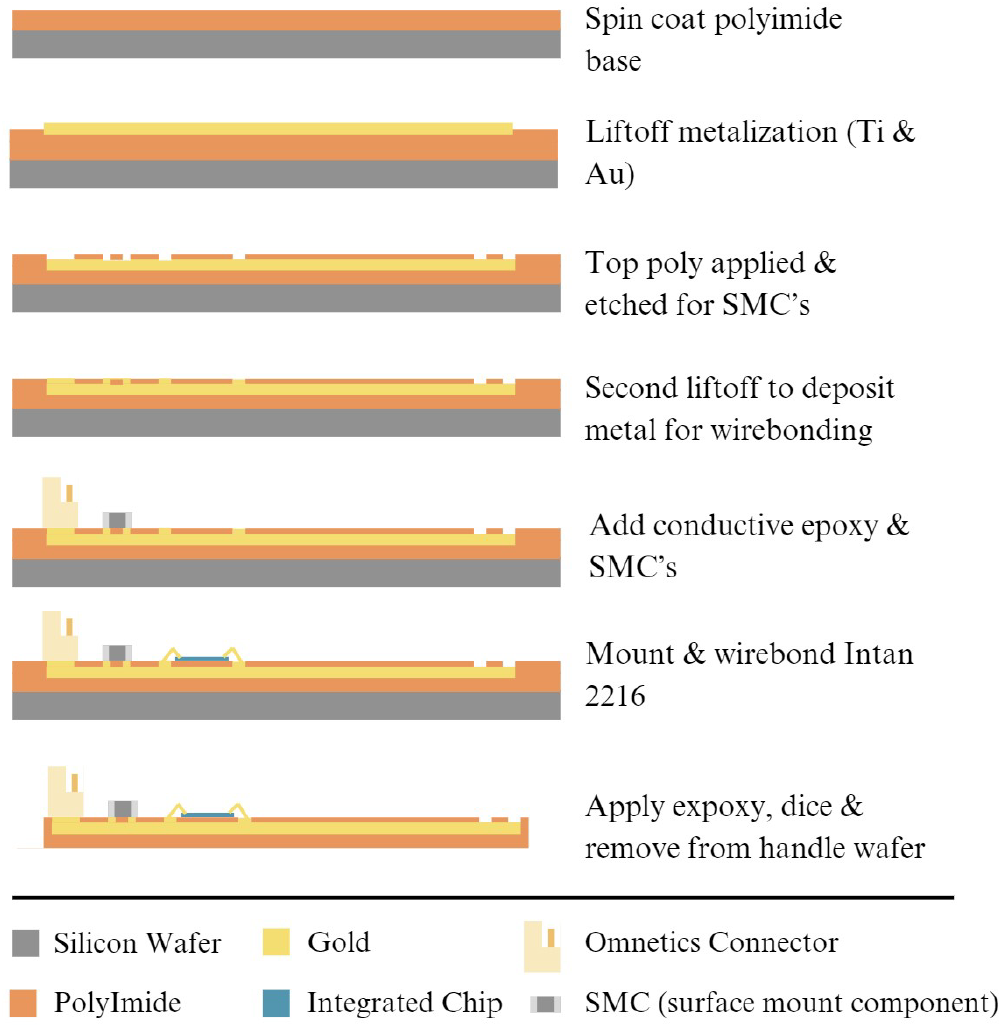
An overview of the process including microfabrication, and benchtop assembly

### C. Benchtop Assembly

To add the surface mount components (SMCs) a stencil was made to “screen print” conductive epoxy on the surface of the FMEA. The conductive epoxy was applied by using a glass slide to press the epoxy through the stencil, and the four resistors (0402, Taiyo Yuden, JPN), two capacitors (0402, Panasonic, JPN), and omnetics connector(A79623, Omnetics Corp, USA) were placed manually under a microscope. Next, the INTAN RHD 2216 is placed using a vacuum pencil and secured using Devcon epoxy (14250, ITW Performance Polymers, USA). Wirebonding using 25µm gold wire was then used to connect the bond pads on the Intan integrated chip (IC) to the flexible substrate. After wirebonding the device is tested by plugging the device into the recording hardware via an SPI cable and ensuring the device is recognized by the software. The SPI cable communicates via 12 necessary connections which include CS±, MOSI ±, MISO1±, MISO2±, VDD, and Ground. Then the device is covered in Devcon epoxy (14250, ITW Performance Polymers, USA) to ensure that the device has rigidity to be removed from the handle wafer. After the epoxy is cured the polyimide array is laser-diced using the Optec Femtosecond laser which creates the shape of the device, and is removed from the handle wafer.

### D. Impedance Testing

After fabrication was completed, we tested the performance by measuring the impedance. For this we connected the device through the SPI cable to an INTAN RHD acquisition board in slot A. Next approximately 8ml of sterile saline solution (0.9% NaCl, AddiPak, USA) was deposited on the electrodes of the device while the device was still on the handle wafer. A ground wire was attached to the acquisition board and the other end was submerged in the saline solution approximately 5mm from the electrodes. At this time, we tested the impedance three times on the 32 differential electrodes. This is a function of the INTAN open access interface (Version 3.3.1) that works with the acquisition board and can be acquired from INTAN’s website [13].

## III. Results

Multiple devices were fabricated to test the feasibility of this method. The features of this design supported our goal of reducing the footprint by 74% and increasing the channel density from 1:1 to 1:3.2. Compared to the previous design requiring 32 connections for 32 electrodes, the new device only requires 10 connections for the same number of electrodes.

Hence, an improvement in channel density. A high-resolution image of the electrodes and the 51 wirebonds is shown in Figure 3.

**Fig. 3.**
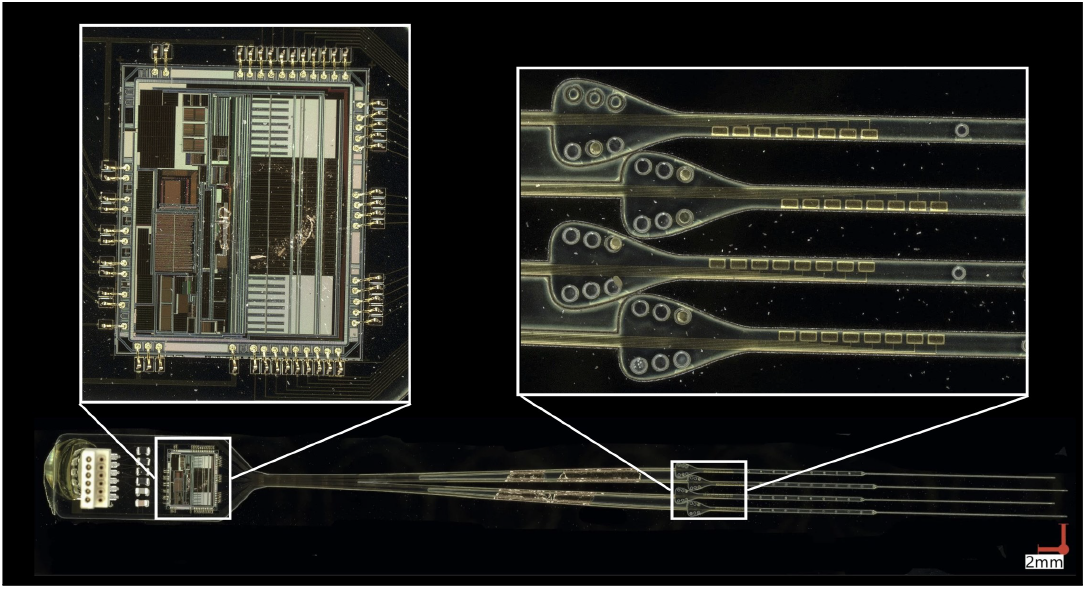
Highlights two features of the device, the 51 wirebonds that connect the bare die to the flexible array, and secondly the electrodes that measure 100µm by 200µm. The tabs on the electrodes, as well as the vias assist with handling the electrodes, and attaching them to a muscle.

Testing the electric characteristics of the device was measured to ensure the recording quality was not affected by the device architecture. The impedance measurements show that the device performed as expected, measuring an average of 100*k*Ω, with a distribution of all electrode impedances shown in Figure 4. These impedances are similar to other gold pad electrodes, and can be improved by adding PEDOT:PSS [14]. Other recent studies show that this level of impedance is acceptable for spike sorting applications [15] and comparing the impedance of this device with non-integrated devices in previous work[1] confirms that we are below the threshold for single motor unit detection. This impedance-threshold is dictated by the host animal’s tissue which is 10 times higher than the impedance of the device.

**Fig. 4.**
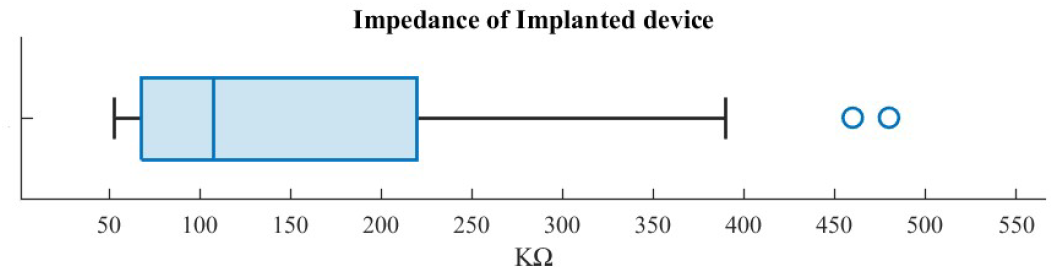
The impedance was measured three times at 1kHz, and each channel recorded a discrete entry.

After electrical characterization, the device was implanted *in-vivo* in an acute setting, on a mouse’s digastric muscle [14]. We chose the digastric muscle due to the amplitude of the spike and the fact that it is active while the mouse is sedated. We gathered data over a 20-minute period that was comparable to measuring with the previous FMEA + INTAN Heastage demonstrated in the Myomatrix arrays. In Figure 5, we show sections of the recordings that have the identifiable motor unit waveform.

**Fig. 5.**
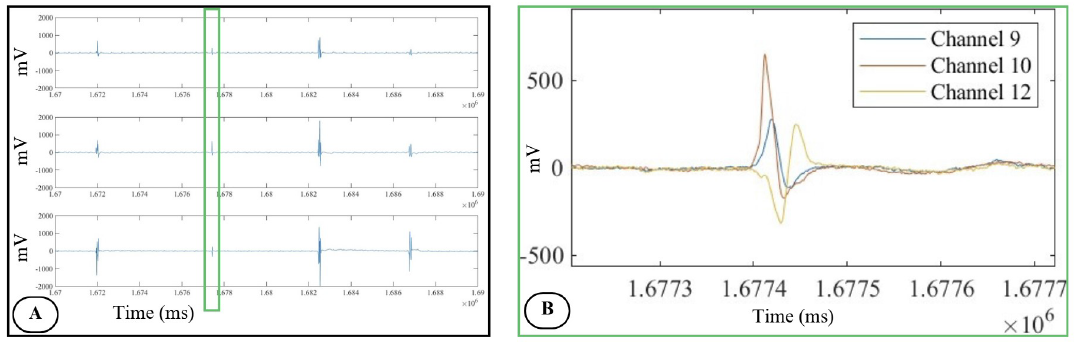
These figures represent the raw EMG data with no filtering applied, plotted by channel in matlab. (A) Higher resolution subset of the channels shown in B, with timing in ms, and the amplitude set to ±2000 mV. (B) Overlays a subset of A showing the change in amplitude across channels 9, 10, and 12 for the same motor unit

## IV. Discussion

### A. Channel density

By establishing a robust method for integrating bare-die chips directly onto Myomatrix / FMEA, we have successfully increased channel density. This device has a channel density of 1:3.2, achieved through the 2216 bare die. The method does not apply to just this chip. With INTAN’s bare die offerings, we can combine RHS (stimulation chips) and higher electrode count RHD (recording chips) bare die to create higher channel density, and control different experiment types. Without changing the method and applying the design changes to the substrate, we can combine multiple RHD2164 bare die to further increase channel density. For smaller animals, we must consider the target muscle, ensuring the device does not become too bulky while increasing channel density. The advantage of this, for EMG specifically, lies in the flexible substrate that can be cut to fit any muscle structure, and the size of the electrodes can be designed accordingly to fit the muscle. With the high SNR capabilities of these devices, these improvements will create a higher-quality data set.

### B. Conclusion

With the Myomatrix’s introduction, we demonstrated a device that can acquire high-resolution EMG and is capable of detecting single motor units. This enabled a new quantitative data set that can be used to study neuromuscular disorders like Parkinson’s and ALS. With this novel device, we have addressed a critical design limitation that created a bottleneck when increasing the number of electrodes. With this new device architecture, we can continue to increase the electrode count and keep a small device footprint. To conclude, this research presents a robust process to reduce the device’s footprint and increase channel density while being applicable to all types of *in-vivo* electrophysiology.

